# Crystal structure of SARS-CoV-2 main protease in complex with a Chinese herb inhibitor shikonin

**DOI:** 10.1101/2020.06.16.155812

**Authors:** Jian Li, Xuelan Zhou, Yan Zhang, Fanglin Zhong, Cheng Lin, Peter J. McCormick, Feng Jiang, Huan Zhou, Qisheng Wang, Jingjing Duan, Jin Zhang

## Abstract

Main protease (M^pro^, also known as 3CLpro) has a major role in the replication of coronavirus life cycle and is one of the most important drug targets for anticoronavirus agents. Here we report the crystal structure of main protease of SARS-CoV-2 bound to a previously identified Chinese herb inhibitor shikonin at 2.45 angstrom resolution. Although the structure revealed here shares similar overall structure with other published structures, there are several key differences which highlight potential features that could be exploited. The catalytic dyad His41-Cys145 undergoes dramatic conformational changes, and the structure reveals an unusual arrangement of oxyanion loop stabilized by the substrate. Binding to shikonin and binding of covalent inhibitors show different binding modes, suggesting a diversity in inhibitor binding. As we learn more about different binding modes and their structure-function relationships, it is probable that we can design more effective and specific drugs with high potency that can serve as effect SARS-CoV-2 anti-viral agents.

---

Severe acute respiratory syndrome coronavirus 2 (SARS-CoV-2), an RNA virus, infects the general population at different ages and can cause severe acute respiratory syndrome in high risk individuals.^1^ The main protease (M^pro^, also known as 3CL^pro^) is essential for the production of infectious virions and play a critical role in the replication of SARS-CoV-2.^2^ M^pro^ is thus an attractive target for the development of drugs against SARS-CoV-2 and other coronavirus infections. M^pro^ of SARS-CoV-2 is a cysteine protease with relatively high sequence homology to other coronavirus main proteases. The catalytic dyad of M^pro^ is formed by His41 and Cys145. A number of studies using either in silico ligand docking or drug discovery based on available structures have been performed to discover new M^pro^ binding agents. Currently, the inhibitors designed for M^pro^ are covalently bound and peptidomimetic, both properties which lend themselves to potential toxicity due to non-specific reactions with host proteins. One previously identified inhibitor was (±)-5,8-dihydroxy-2-(1-hydroxy-4-methyl-3-pentenyl)-1,4 naphthoquinone, termed shikonin, which derives a Chinese herb, and is a major active component of the roots of *Lithospermum erythrorhizon.*^4^

To find new scaffold and non-covalent inhibitors and reveal further details of inhibitor binding, we determined the crystal structure of M^pro^ in complex with shikonin. The structure reveals a novel binding mode that opens new opportunities for future drug development targeting the M^pro^ protease (Fig. 1a, b). Shikonin and its derivatives have been reported to have antiviral, antibacterial, anti-inflammatory and anti-tumor effects.^4,5^ Previous data have shown shikonin has 15.75 μM IC_50_ to M^pro^ protease.^5^ These data, in combination with the structure revealed in this study highlight shikonin as a starting point for developing future novel non-covalent antiviral molecules.

**Fig. 1.**
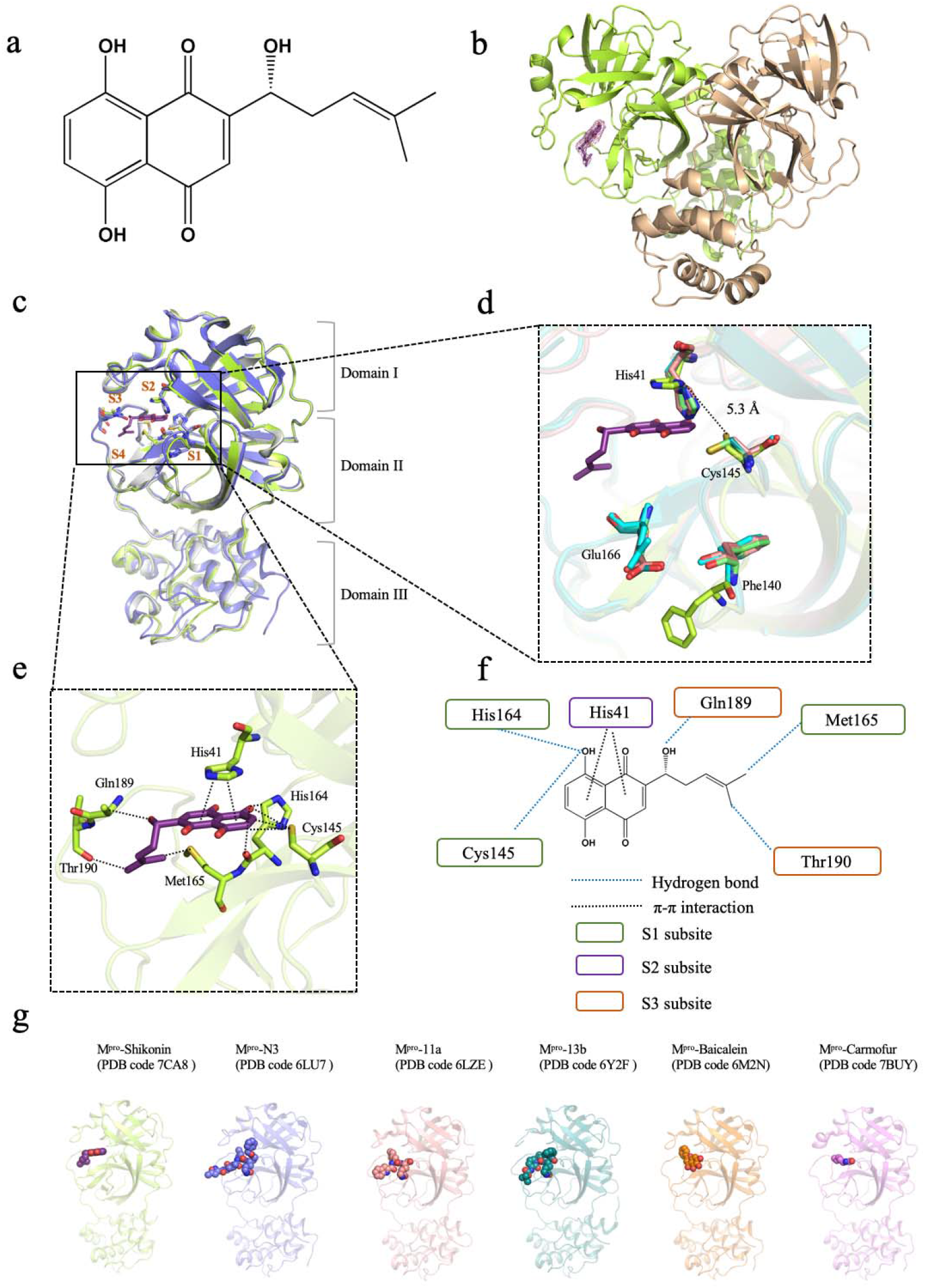
Crystal structure of SARS-CoV-2 main protease (M^pro^) in complex with a Chinese herb inhibitor shikonin. a Chemical structure of the non-colavent inhibitor shikonin. **b** Structure of M^pro^ dimer. One protomer of the dimer with inhibitor shikonin is shown in green, the other is shown in wheat. The contour level is at 1σ. **c** Comparison of SARS-CoV-2 M^pro^ structures. Brown symbols S1, S2, S3 and S4 indicate the substrate binding pockets. Structure of ^Shi^M^pro^ is shown in green. Structure of M^pro^ with N3 is shown in light blue. Structure of ^apo^M^pro^ is shown in grey Carbon atoms of shikonin are magenta, oxygen atoms are red. Hydrogen bonds and π-π interactions are indicated by dashed black lines. **d** Conformational difference in catalytic site His41-Cys145. Residues of M^pro^ structure with shikonin are shown in green. **e** A zoomed view of shikonin binding pocket. **f** Schematic interaction between shikonin and M^pro^. Hydrogen bonds and π-π stacking interactions are shown as blue dashed lines and black dashed lines, respectively. Green circle indicates conserved residues in S1 subsite. Purple circle indicates conserved residues in S2 subsite. Orange circle indicates conserved residues in S3 subsite. **g** Crystal structures of M^pro^-inhibitor complexes from previously reported structures presenting diverse inhibitor-binding sites. M^pro^ structures are shown in cartoon representation and the inhibitors are shown as sphere models with transparent surfaces. The representative structures of M^pro^ along with covalent inhibitors, N3 (PDB code 6LU7), 11a (PDB code 6LZE) and 13b (PDB code 6Y2F) are shown. Similarly, structures for M^pro^ bound to natural products shikonin (PDB code 7CA8) and baicalein (PDB code 6M2N), and antineoplastic drug carmofur (PDB code 7BUY) are shown.

## Overall structure

The crystal structure of M^pro^ in complex with shikonin (^Shi^M^pro^) has been determined at 2.45 Å resolution using a previously described M^pro^ construct (Table 1, Fig. 2a). ^Shi^M^pro^ structure shows the same overall fold observed for the previous apo state structure of M^pro^ at pH 7.5 (^apo^M^pro^).^3^ The r.m.s. difference in equivalent Cα positions between apo and ^Shi^M^pro^ is roughly 0.3 Å for all the residues (Fig. 1c). Some key residues in the oxyanion hole and N-finger were found to be disordered in the apo state structure. These residues are located near the active site of the enzyme and therefore participate in the binding of substrates or inhibitors. However, unlike the apo state of M^pro^ at pH 7.5, the current structure contains residues with clear density for both protomers of the protease in the crystal. Interestingly, shikonin binds to protomer A but not to protomer B (Fig. 2b, c). The reasons are unclear but may be due to the relatively low affinity of shikonin.^5^ Structurally, the two monomers have essentially similar conformation with slight difference in oxyanion loop (Fig. 3). There are, however, remarkable differences both in the conformation of the protease monomer between this inhibitor-bound complex and the apo state reported earlier by us.^3^ Residues in the oxyanion loop became more ordered due to inhibitor binding, as residues 140-145 appear to interact with the inhibitor. Notably, an unprecedented conformational difference in the catalytic dyad His41 and Cys145 is observed, leading to a steric clash between the previously reported inhibitors and His41-Cys145 catalytic site (Fig. 1d).^5–9^

**Table1.** Statistics for data processing and model refinement of main protease of SARS-CoV-2 in complex with shikonin.

|  |  |
| --- | --- |
| PDB code | 7CA8 |
| Synchrotron | SSRF |
| Beam line | BL17U1 |
| Wavelength (Å) | 0.97918 |
| Space group | P2<sub>1</sub>2<sub>1</sub>2<sub>1</sub> |
| *a*, *b*, *c* (Å) | 67.96, 102.37, 103.54 |
| *α*, *β*, *γ* (°) | 90.00, 90.00, 90.00 |
| Total reflections | 356383 |
| Unique reflections | 27221 |
| Resolution (Å) | 2.45(2.51-2.45) |
| R-merge (%) | 13.4(163) |
| Mean *I* / *σ* (*I*) | 15.7/2.2 |
| Completeness (%) | 99.7(99.5) |
| Redundancy | 13.1(13.7) |
| Resolution (Å) | 56.81-2.45 |
| *R*<sub>work</sub>/*R*<sub>free</sub>(%) | 22.24/27.62 |
| Atoms | 4917 |
| Mean temperature factor (Å<sup>2</sup>) | 56.9 |
| Bond lengths (Å) | 0.008 |
| Bond angles (°) | 1.019 |

Values in parentheses are for the highest-resolution shell.

**Fig. 2.**
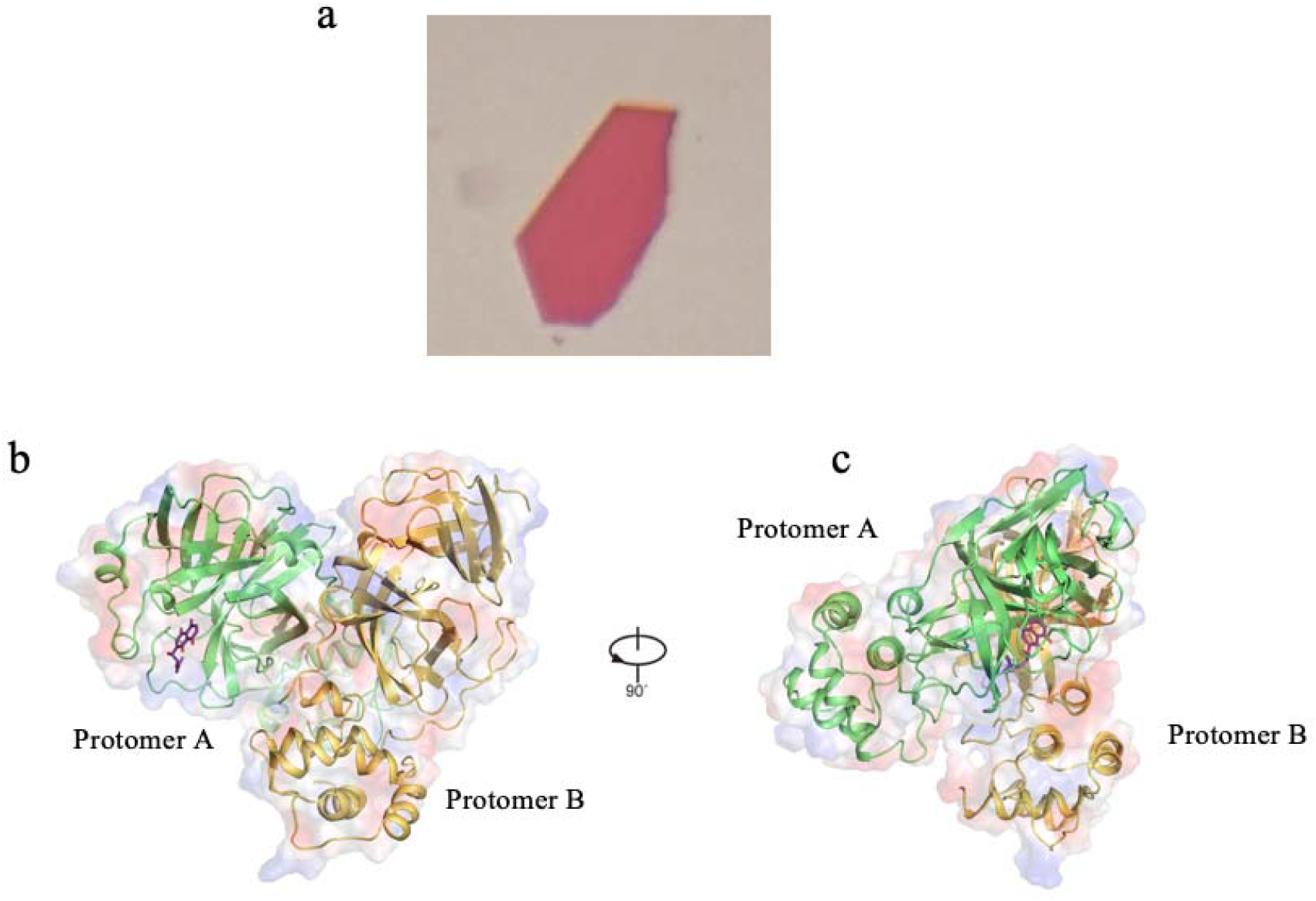
Overall structure of main protease (M^pro^) of SARS-CoV-2 in complex with shikonin. **a** M^pro^ crystal soaked with shikonin. **b,c** Different views of the homodimer of M^pro^. Protomer A is in green, protomer B is in wheat, shikonin is presented as purple sticks.

**Fig. 3.**
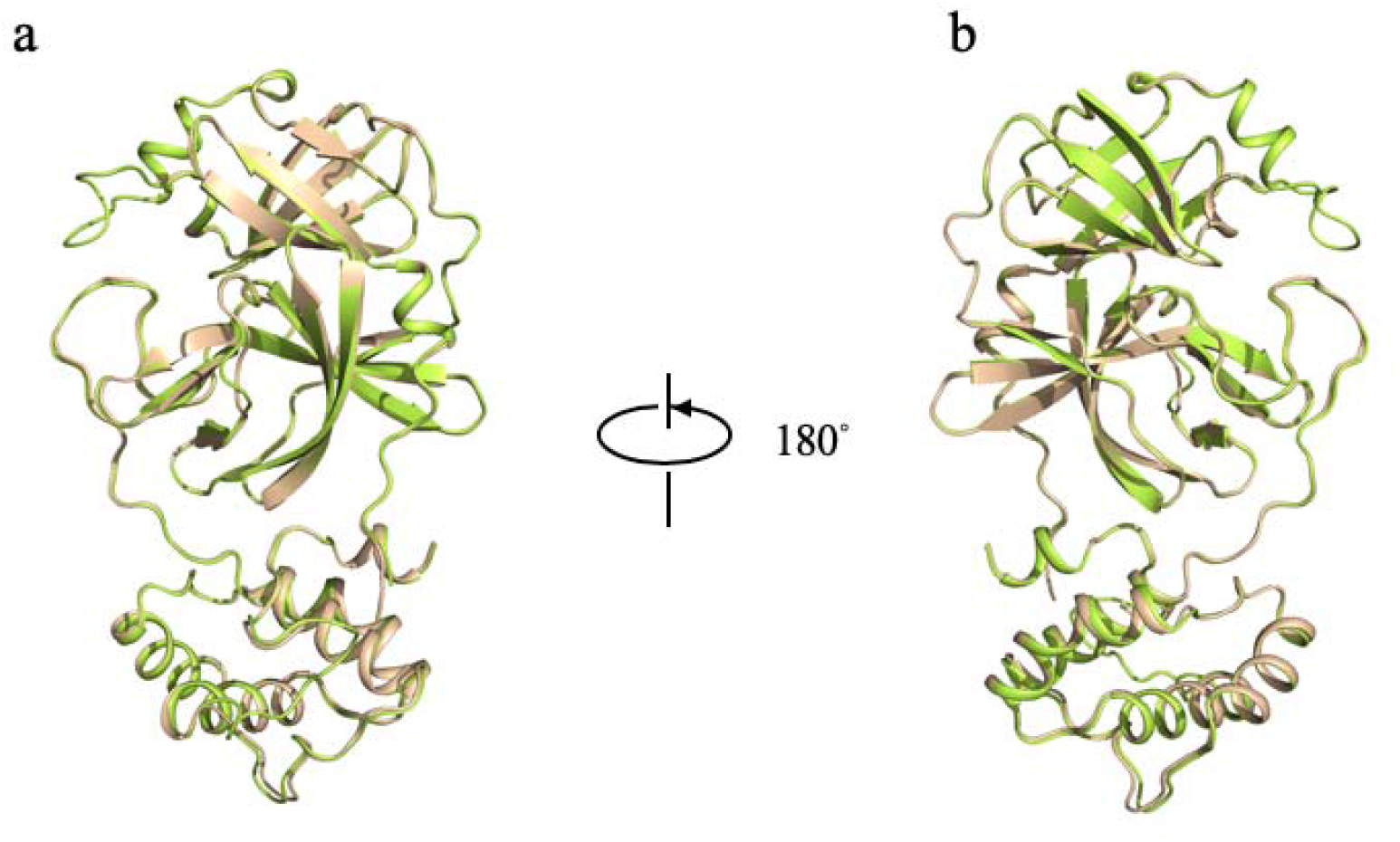
Structural comparison of the protomer A (green) of the ^Shi^M^pro^ with protomer B (wheat).

### Unique binding mode of the non-covalent inhibitor shikonin

An overlay of the ^Shi^M^pro^ structure with previously solved inhibitor-bound structures shows relatively high spatial conservation of the three domains (Fig. 1c, Fig. 4). Shikonin contains 1,4-naphthoquinone (1,4-NQ) that consists of a benzene moiety and a fully conjugated cyclic diketone with the carbonyls are arranged in para position, referred to as the inhibitor head group, and a chiral six-carbon side-chain with the hydroxy group at C-1, defined as the ligand tail. The inhibitor binding pocket can be described as a narrow cleft surrounded by S1-S4 subsites (Fig. 1c). Shikonin establishes a hydrogen bond network with the protease polar triad Cys145, His164 and Met165 located on the S1 subsite. The aromatic head groups of shikonin forms a π-π interaction with His41 on the S2 subsite. The hydroxy and methyl group of the isohexenyl side chain tail has two H-bonding with Gln189 and Thr190 on the S3 subsite, respectively. Superimposing ^Shi^M^pro^ with other inhibitor-bound structures reveals striking difference in the arrangement of the catalytic dyad His41-Cys145 and smaller, but substantial, differences in Phe140 and Glu166. In both covalent inhibitors-bound structures, the inhibitor binds to the Sγ atom of Cys145. In the case of current structure with shikonin, the side chain of Cys145 adopts a different configuration, forming hydrogen bond with shikonin (Fig. 1d, e). In addition, the imidazole group of His41 pointing towards the binding pocket in other structures flip outward, opening a way for the entry of shikonin. The distances between His41 N_∊_2 and Cys145 Sγ are 5.3 Å in ^Shi^M^pro^ structure, and the distance is significantly longer than that observed in any other main protease of reported structures (Fig. 1d).^5–9^ Phe140 no longer has π-π interaction with His163, and the phenyl ring of Phe140 undergoes dramatic conformational change and moves outward to the solvent (Fig. 1d). The side chain of Glu166 is flexible and adopts an inactive conformation in both apo and ^Shi^M^pro^ structures, but is well ordered in the other known structures (Fig. 1d). It has been shown that Glu166 is critical in keeping the substrate binding pocket in a suitable shape by forming hydrogen bond with peptidomimetic inhibitors and N terminal residue in the other protomer.^8^ This may explain why Glu166 is strictly conserved among all main proteases.

**Fig. 4.**
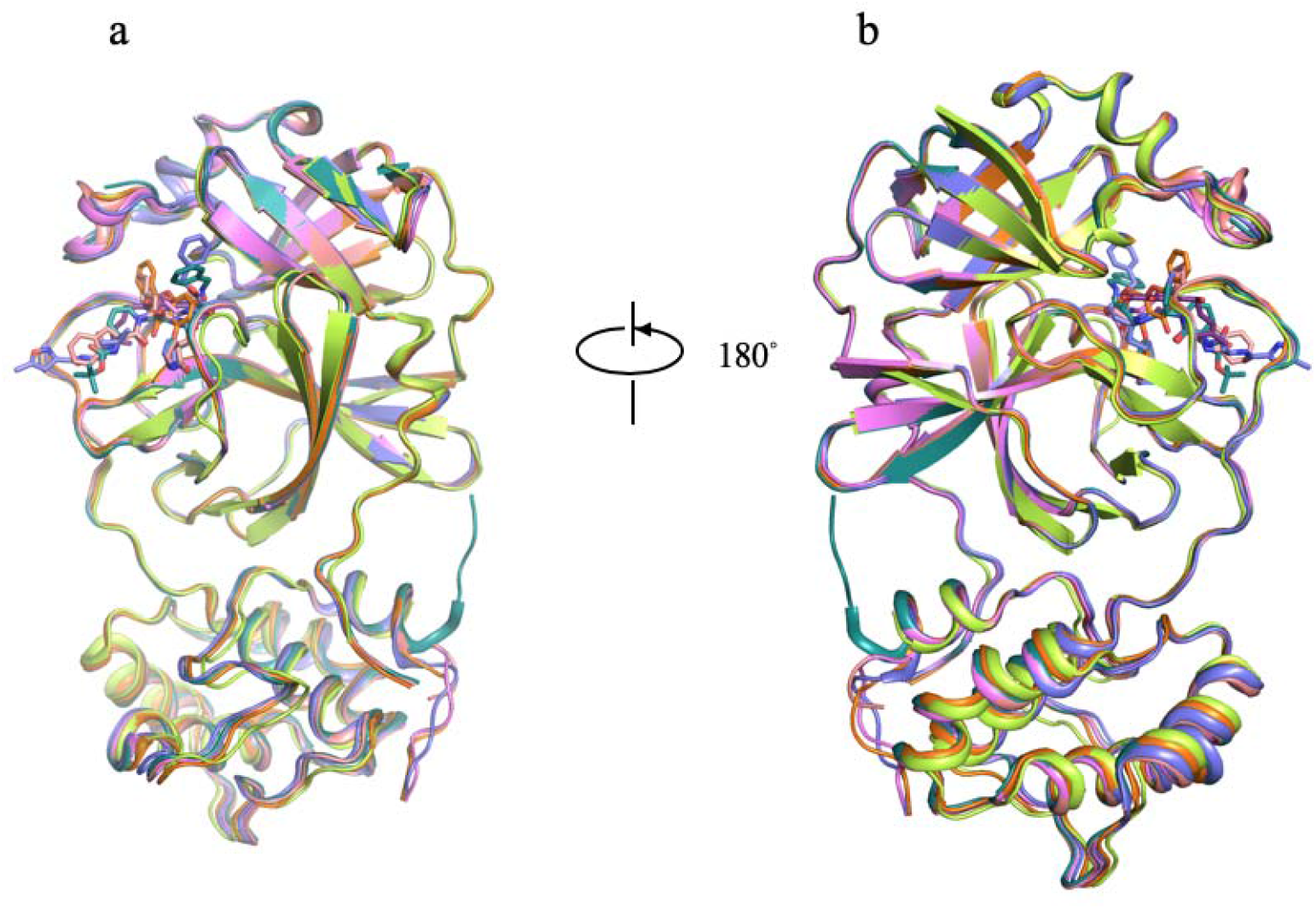
Superposition of all known inhibitor-bound structures for M^pro^ of SARS-CoV-2.

### Two water molecules in the active site of the protease

The ^apo^M^pro^ structure reveals the presence of two water molecules in the substrate binding site (Fig. 5a). Water 1 forms hydrogen bond network involving Phe140, His163 and Glu166 located in the S1 pocket, stabilizing the oxyanion hole in the apo state structure.^3^ Another water molecular (water 2) hydrogen bonded with His41 and Cys145 in the apo state structure is occupied by the shikonin in the ^Shi^M^pro^ structure (Fig. 5b). However, these two water molecules are not observed in the ^Shi^M^pro^ structure. We propose that inhibitors that are able to replace these water molecules may have significant improvement of potency for the protease, as was observed when the two water molecules in the substrate binding pocket of M^pro^ are replaced by the inhibitors (Fig. 5c).^5–9^

**Fig. 5.**
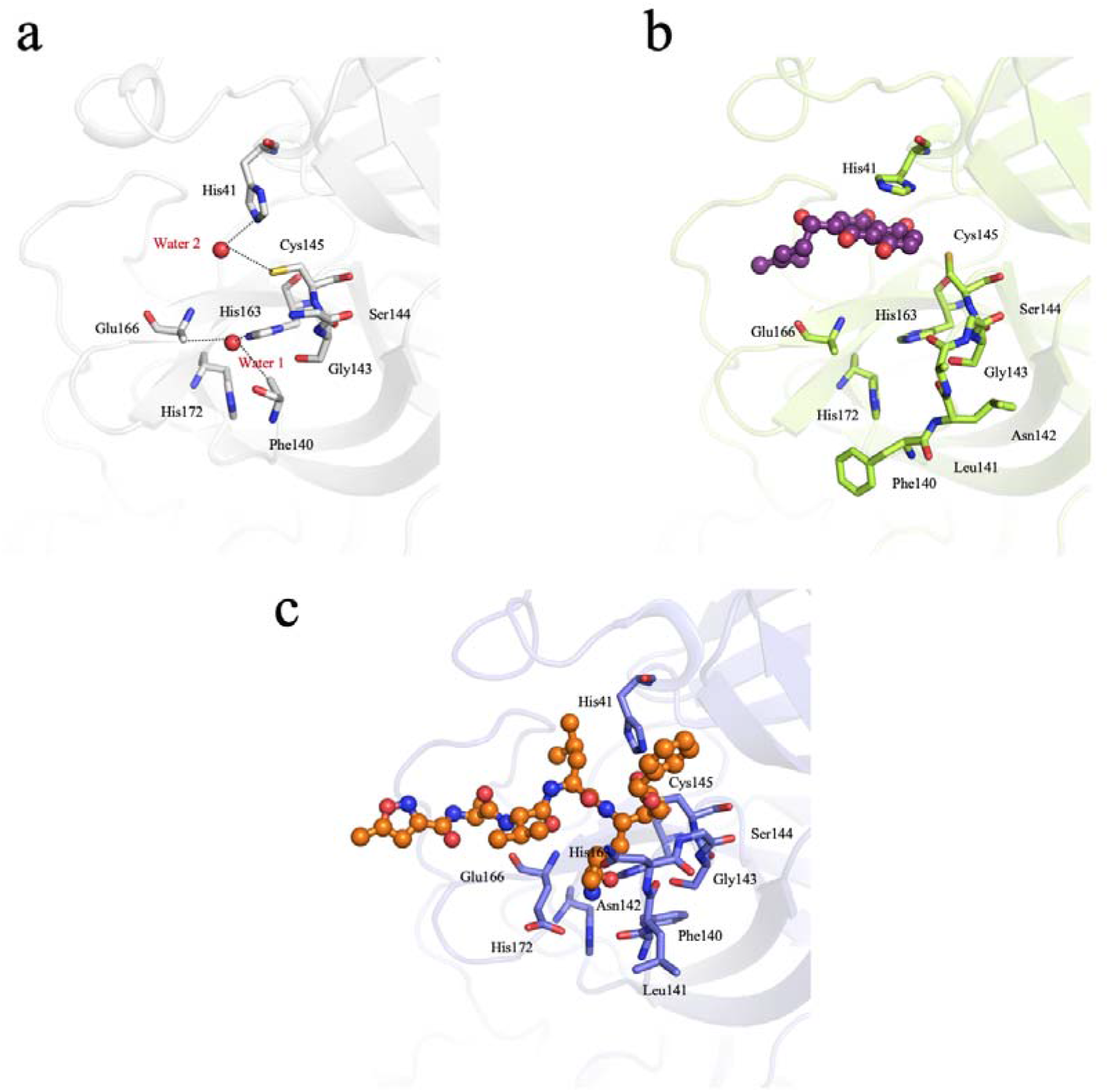
Inhibitor binding mode to M^pro^ of SARS-CoV-2. The inhibitor binding residues in ^apo^M^pro^, ^Shi^M^pro^ and M^pro^ with N3 are shown in grey, green and light blue, respectively. Water molecules are shown in red sphere.

## Conclusion

The current global pandemic has increased the urgency for novel small molecule inhibitors to slow or block SARS-CoV-2 viral propagation. Here we have shown that the napthoquinone, shikonin, binds in a unique mode to the M^pro^ protease. Our structure reveals three novel interactions in the substrate/inhibitor binding pocket, 1) the π stacking interaction between the shikonin naphthoquinone core and side chain of His41 from the S2 subsite, 2) hydrogen bonds with Cys145, His164 and Met165 in the S1 pocket and 3) hydrogen bonds with Gln189 and Thr190 in the S3 pocket (Fig. 1e, f). To date it has been shown that the covalent and peptidomimetic inhibitors identified bind to S1/S2/S4 site, while camfour only binds to the S2 subsite and another natural product baicalein binds to the S1/S2 pocket (Fig. 1g).^5–9^ The ^Shi^M^pro^ structure highlights a new binding mode of non-covalent and natural product inhibitors, with distinct local conformational changes in the substrate binding pocket, and represents an exciting novel scaffold derived from a Chinese medicinal herb. The M^pro^ structure identified here in complex with natural product shikonin provides an invaluable resource to design improved antiviral drugs for this important therapeutic target.

## Materials and Methods

### Protein purification and crystallization

The cDNA of full-length SARS-Cov-2 main protease 3CL (NC_045512) was optimized, synthesized (Generay, China) and cloned into the pET28a vector. The plasmid was transformed into competent cell E.coli Rosetta DE3. The bacteria were grown in 800mL of LB (Luria-Bertani) broth at 37 °C. When the OD600 reach 0.6-0.8, 500 μM IPTG was added to induce the E. coli expression and then incubated 3-5h at 30°C. The cells were centrifuged at 10000 g for 15min at 4 °C, the supernatant discarded, and the precipitate collected. Buffer A (100 mM Tris/HCl buffer, pH7.5, 300 mM NaCl 10mM imidazole and 5% glycerol) was added to resuspend the collected cells, which were broken up by a JNBIO 3000 plus (JNBI). The supernatant containing the protein was acquired by centrifugation at 30000 g, 4 °C for 3 0min. The supernatant was transferred into a 5 ml Ni-NTA(Ni2+-nitrilotriacetate)column (GE healthcare) and the protein wanted was loaded onto the column. Buffer B(100 mM Tris/HCl buffer)pH 7.5, 300 mM NaCl, 100 mM imidazole, and 5% glycerol) was added into beads as a 30 times column wash. The His tagged protein was eluted by buffer C (50 mM Tris-HCl pH 7.5, 300 mM NaCl and 300 mM imidazole). A Superdex 200 PG gel filtration column (GE healthcare) was used to purify the protein and remove the imidazole, and the buffer changed to buffer C(25 mM HEPES buffer, pH7.5, 30 0mM NaCl, 2 mM DTT and 5% glycerol). Fractions were collected and analyzed by SDS-PAGE. Positive fractions containing the protein were flash-frozen in liquid nitrogen and stored at −80 °C.

The protein was thawed and concentrate at 10 mg/ml in Amicon Ultra-15,10000Mr cut-off centrifugal concentrator (Millipore). 10 mM shikonin was added to the protein in a ratio of 5:1 and bond at 4°C for 2h. The hanging drop vapor diffusion method was useful to gain crystals at 20°C. Under the conditions of crystal described before (0.1M HEPES 7.5, 10% propanol and 20% PEG4000 in 2-3 days). 3

### Data collection, structure determination and refinement

The crystals were tailored with cryo-loop (Hampton research, America) and then flash-frozen in liquid nitrogen to collect better X-ray data. The data set was collected at 100 K on a macromolecular crystallography beamline17U1 (BL17U1) at Shanghai Synchrotron Radiation Facility (SSRF, Shanghai, China). All collected data were handled by the HKL 2000 software package. The structure was determined by molecular replacement with PHENIX software. The of 7C2Q was referred as a model. The program Coot was used to rebuild the initial model. The models were refined to resolution limit 2.45 Å by using the PHENIX software. The complete wanted data collection and statistics of refinement are shown in Table 1. The structure has been deposited in PDB (PDB code 7CA8).

## Acknowledgments

We thank the SSRF BL17U1 beam line for data collection and processing. Jian Li was supported by the Open Project of Key Laboratory of Prevention and treatment of cardiovascular and cerebrovascular diseases, Ministry of Education (No. XN201904), Gannan Medical University (QD201910) and Jiangxi “Double Thousand Plan”. Jin Zhang was supported by the Thousand Young Talents Program of China, the National Natural Science Foundation of China (grant no. 31770795; grant no. 81974514), and the Jiangxi Province Natural Science Foundation (grant no. 20181ACB20014). Peter J. McCormick was supported by the Foreign Talent project of Jiangxi Province. Feng Jiang was supported by the National Natural Science Foundation of China (Nos. 21961003), Natural Science Foundation of Jiangxi Province (Nos. 20192BAB205114), Talent project of Jiangxi Province. Jingjing Duan was supported by the Natural Science Foundation of China (grant no. 31971043). This work was also supported by Ganzhou COVID-19 Emergency Research Project.

## Author contributions

J.L., X.Z., Y.Z., F.Z., and C.L. made constructs for expression and determined the conditions used to enhance protein stability. H.Z., and Q.W. carried out X-ray experiments, including data acquisition and processing. J.L. built the atomic model. J. Z., J.L., Y.Z., P.J.M., and J.D. drafted the manuscript. F.J. contributed to structure analysis/interpretation and manuscript revision. J.Z and J.L. supervised the research.

